# Bat trypanosomatids (first report of *T. wauwau*) in Triângulo Mineiro, Brazil

**DOI:** 10.1101/347146

**Authors:** Pablo de Oliveira Pegorari, César Gómez-Hernández, Cecilia G. Barbosa, Karine Rezende-Oliveira, André Luiz Pedrosa, Juan David Ramirez, Luis E. Ramirez

## Abstract

In this study, trypanosomatids commonly found in bats, including *Trypanosoma cruzi marinkellei, T. dionisii,* and *Leishmania braziliensis,* were identified. Additionally, *T. wauwau* was identified in one specimen of *Anoura caudifer,* and represents the first report of this parasite from the Central West region of Brazil. *T. wauwau* was previously identified by other researchers in the North of the country, in only three species of bats in the genus *Pteronotus: P. parnellii* (Pará and Rondônia states), and *P. personatus* and *P. gymnonotus* (Rondônia). The identification of *T. wauwau* indicates how different trypanosomatids are able to adapt to new host species of bats. This is owing to bats’ high mobility, wide geographic distribution, social behavior, and ability to coexist in large colonies. These characteristics may facilitate the transmission of infectious agents in nature, which are responsible for outbreaks of some zoonoses. Therefore, health authorities should focus on both vertebrates and vectors associated with the environments where these bats are found.

**Author summary:** The prevalence of *Trypanosoma* in bats is high, with *T. cruzi, T. cruzi marinkellei*, and *T. dionisii* as the most prevalent infective species. This study reports for the first time the presence of *T. wauwau* in the southeast region of Brazil in the bat *Anoura caudifer.* Although this species of *Trypanosoma* has been found in bats of the genus *Pteronotus*, it was not detected in any other genus, including in the bats that share the same shelter with *Pteronotus*. The species *T. wauwau* was found infecting bats only in Brazil. Its occurrence was restricted to the northern region of the country, in the states of Pará, infecting the species *P. parnellii* and in *Rondônia* infecting *P. personatus, P. gymnonotus* as well as *P. parnellii*. Although its morphology is similar to that of *T. cruzi*, little is known about the development of *T. wauwau*, both in its vertebrate host and the existence of a plausible invertebrate vector. Its characteristics include its inability to develop in mammalian cells and its non-infectiousness in mice and triatomine insects. Further research, through molecular studies, may provide important and valuable data for understanding the origin, evolution, and global distribution of, and the association between the different species of *Trypanosoma* and their hosts.

## Introduction

Until now, more than up to 30 species of trypanosomatids have been isolated from bats [1–4]. The most frequently reported species of *Trypanosoma* in bats are *T. cruzi, T. cruzi marinkellei, T. dionisii, T. hedricki, T. myoti, T. leonidasdeanei, T. desterrensis, T. pifanoi, T. pessoai, T. megaderma*-like, *T. theileri*, and *T. rangeli* [5–10]. An additional species, *T. wauwau*, was also recently described in Brazil [9–10]. Another group of bat-related trypanosomatids are those in the genus *Leishmania*, with L. *infantum chagasi* reported infecting bats in Venezuela [11], later, *L. mexicana* was found in the bats in southeastern Mexico [ 12] and the species *L. amazonensis, L. braziliensis,* and *L. infantum chagasi* were found in Brazil[3, 13–16]. Because of the detection of such medically important trypanosomatids in bats, epidemiological surveillance research is necessary in regions with endemic and non-endemic disease-causing species that may pose a risk of human infection.

Given the above observations and concerns, the objective of the present study was to evaluate the presence and identity of the possible trypanosomatid species (*Trypanosoma* spp. and *Leishmania* spp.) in bats of the Triângulo Mineiro Region of Brazil.

## Material and methods

### Area of study and capture of bats

This study was carried out in the city of Ituiutaba, Minas Gerais (MG), Brazil, located in the Triângulo Mineiro mesoregion (lat 18°58‘08’ S, lon 49°27‘54’ W; altitude: 544 m; area: 2595.2 km^2^), west of MG, Brazil.

Bats were captured from November 2014 to September 2015, at night between 18h00 and midnight (0h00) using mist nets, and during the day in shelters using manual nets. The identification of bats species was done based on taxonomic keys to the family [ 17], genus, and species levels [ 18–19].

### Blood collection

A total of 216 bats were collected, and 0.5 to 1.0 mL of blood was collected from each specimen by cardiac puncture. Of these samples, 25 µL was used to estimate microhematocrit, while the remaining blood was stored in EDTA V/V and guanidine solution (6 M Guanidine–HCl and 0.2 M disodium) at 4°C until further use.

### Identification of trypanosomatid species

DNA extraction was performed using a GeneJET Genomic DNA Purification^®^ kit from Thermo Scientific, according to the manufacturer’s instructions.

For detecting *Leishmania* DNA, the primers HSP70F and HSP70R (5′CCGCCCATGCTCTGGTACATC 3′) were used, whose target is the HSP70 gene of *Leshmania* spp. [20]. For detecting *Trypanosoma* DNA, a Nested PCR was performed, in which the primers used in the first reaction were TRY927F (5’-GAAACAAGAAACACGGGAG-3’) and TRY927R (5’- CTACTGGGCAGCTTGGA-3’), and those used in the second reaction were SSU561F (5’- TGGGATAACAAAGGAGCA-3’) and SSU561R (5’- CTGAGACTGTAACCTCAAAGC- 3’), whose target was the 18S rDNA region [21]. Electrophoresis was performed on 6% polyacrylamide gel, stained with silver. Sequencing was performed at ACTGene Análises Moleculares (Brazil). Chromatograms were analyzed using ChromasPro 2.1.4, and consensus sequences were generated. The Phred threshold value was set at > 20. Sequences were aligned using SeaView 4.5.2, with the Muscle algorithm. Maximum Likelihood trees using Neighbor-Joining methods were generated using MEGA7: Molecular Evolutionary Genetics Analysis version 7.0 for bigger datasets [22], with 1000 bootstrap iterations, and including reference sequences from trypanosomatids retrieved from GenBank.

### Statistical analyses

The Chi-square test was used to assess the possible association between the dietary habits of bat hosts and the positivity of their PCR tests for both *Leishmania* and *Trypanosoma*, using the TIBCO^®^ Statistica™ program [23], with a significance level of 5% (P < 0.05).

### Ethical considerations

Bats were captured and manipulated according to the recommendations of the Brazilian Institute of Environment, with authorization for activities with scientific purpose number 45132-1 (IBAMA-SISBIO). The procedures used were approved by the Animal Research Committee of the Federal University of Triângulo Mineiro, as per protocol No. 51.

## Results

### Identification of *Trypanosoma* species

The captured animals (216) belonged to nine different bat species. Of these, 43 (19.90%) presented positive microhematocrit results for trypanosomatids. Among the blood samples analyzed, 18 (8.33%) were found positive for *Trypanosoma* spp. DNA, most (77.77%) of which came from nectarivorous bats (Table 1). However, according to the results of the Chi-square test, there was no significant statistical association between the dietary habits of hosts and infection with *Trypanosoma* spp. (p = 0.205) or *Leishmania* spp. (p = 0.635).

**Table 1.**
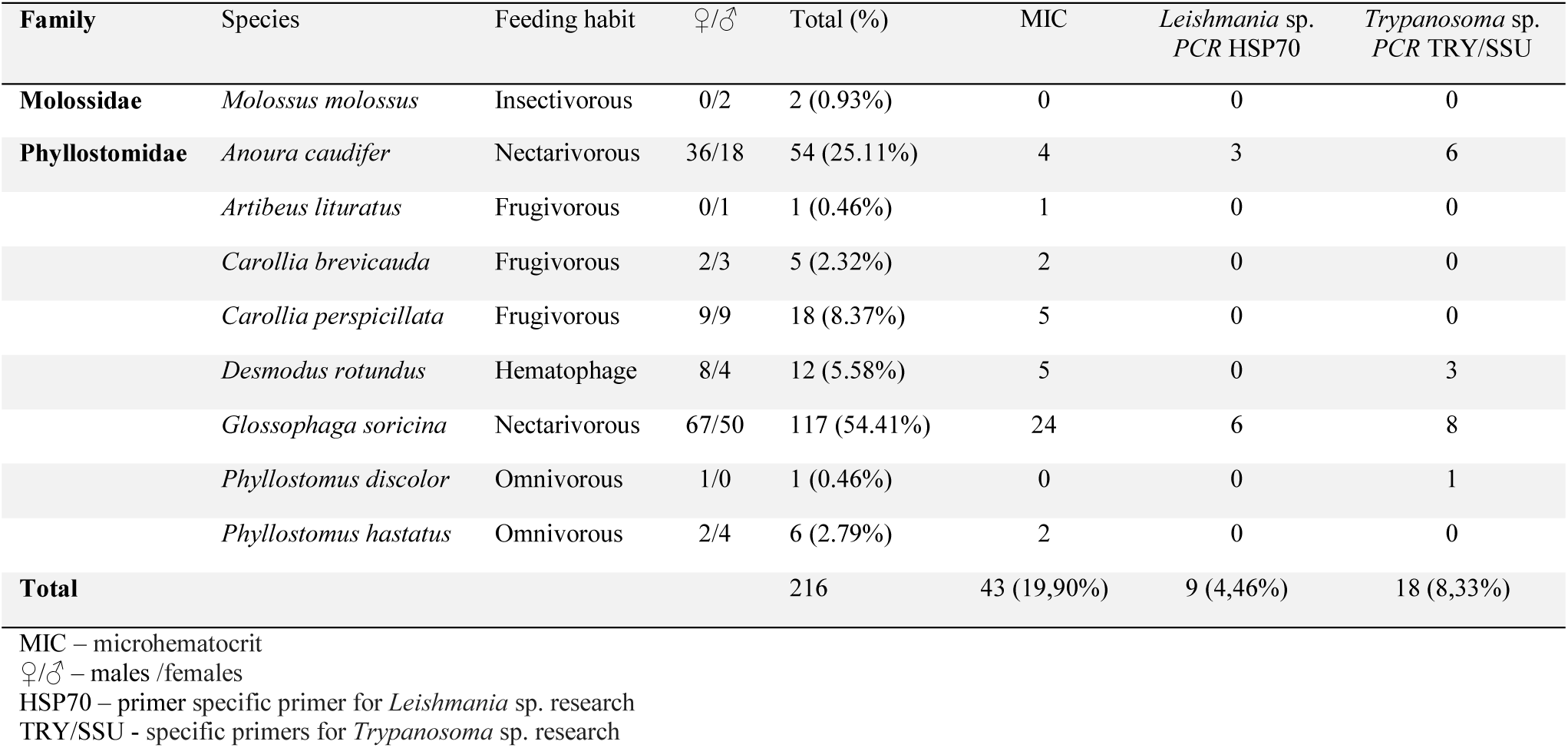
Bat species captured in Minas Gerais, Brazil by family, feeding habits, sex and positivity.

The species of bats with the highest infection rates were *Desmodus rotundus, Anoura caudifer,* and *Glossophaga soricina* (25.00%, 11.11%, and 6.83%, respectively). The species of trypanosomes identified were *T. cruzi marinkellei* and *T. dionisii* in various bat species, and *Trypanosoma wauwau* in *Anoura caudifer* (Table 2, Figure 1).

**Figure 1.**
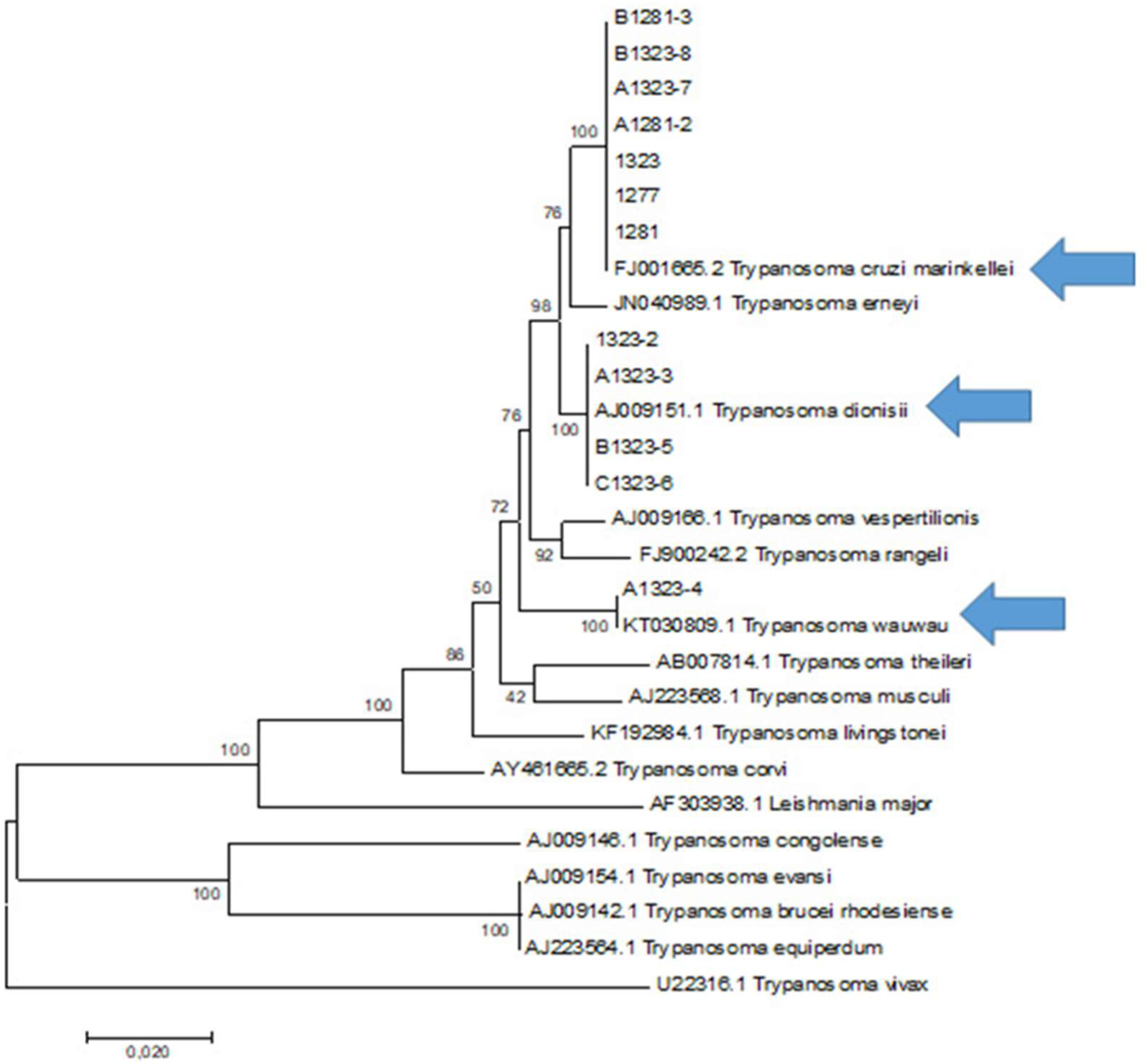
Phylogenetic analysis by the Maximum Likelihood method. The phylogenetic analysis was performed using the Maximum Likelihood method based on the Tamura-Nei model with 1000 bootstraps. A total of 1954 bases were analyzed together. All positions containing gap were eliminated. The phylogenetic analyzes were performed in the MEGA7 program. The arrows indicate the species found in the sequencing of the present study.

**Table 2.** Trypanosomatids isolates, isolate code, bat hosts and primers used to DNA sequencing.

| Trypanosomatids species | Quantity | Individual isolate code | Host | Primer used |
| --- | --- | --- | --- | --- |
| *Leishmania braziliensis* | 2 | Mo205Bac/ Mo224Bac | *Anoura caudifera* | HSP70Leish |
| *Leishmania braziliensis* | 1 | Mo110Bac | *Phyllostomus discolor* | HSP70Leish |
| *T. cruzi marinkellei* | 4 | Mo220ID/ Mo254/ Mo256/ Mo258 | *Glossophaga soricina* | TRY927/SSU561 |
| *T. cruzi marinkellei* | 1 | Mo225 | *Desmodus rotundus* | TRY927/SSU561 |
| *T. cruzi marinkellei* | 1 | Mo261 | *Anoura caudifera* | TRY927/SSU561 |
| *Trypanosoma dionisii* | 2 | Mo176/ Mo218 | *Anoura caudifera* | TRY927/SSU561 |
| *Trypanosoma wauwau* | 1 | Mo184 | *Anoura caudifera* | TRY927/SSU561 |

### Identification of *Leishmania* species

Of the samples analyzed, nine (4.46%) were positive for *Leishmania* spp. (Table 1). The only species of *Leishmania* identified was *L. braziliensis*, in the bats *Anoura caudifer* and *Phyllostomus discolor* (Table 2).

## Discussion

Some species of trypanosomatids have already been found to naturally infect bats [4, 7–10, 24–27]. Many of these trypanosomatids are responsible for important zoonoses, but most of those found in bats remain poorly known. In this work, we investigated the presence of *Leishmania* spp. and *Trypanosoma* spp. in bats from the city of Ituiutaba, Triângulo Mineiro, Brazil, an area previously showing endemism for Chagas disease.

In this study, the use of the direct parasitological method of microhematocrit revealed frequent positive results for trypanosomatids, as has been demonstrated by other studies [4, 28]. It is worth highlighting that the microhematocrit method, in all the field studies carried out by our group, especially in small animals, has shown greater sensitivity or, in certain cases, similar behavior to that of traditional blood culture methods [29].

Regarding PCR positivity, the prevalence of trypanosomatids was similar to that observed by other studies in Brazil [3–4], as was the result that in terms of dietary habits most of these bats were nectarivores. It was also demonstrated in this study that the two most common species that circulate in bats in this region are *T. c. marinkellei* and *T. dionisii,* as has been previously demonstrated [4, 6, 30].

The only species of *Leishmania* found was *L. braziliensis*, in *Anoura caudifer* and *Phyllostomus discolor*. This is the first report of this species of *Leishmania* in these species of bats. In this region, the species *L. infantum, L. amazonensis,* and *L. braziliensis* were previously identified from the bat species *G. soricina* and *M. molossus* [3].

In this study, it was possible to identify *T. wauwau* in only one specimen of *Anoura caudifer* (a nectarivore of the family Phylostomatidae), with this being the first report of this parasite in the Central-West region of Brazil, and in a different species of bat from those previously reported to harbor it. This *Trypanosoma* was recently identified in Brazil in only three species of bats, all in the genus *Pteronotus: P. parnellii* in Pará and Rondônia states, and *P. personatus* and *P. gymnonotus* in Rondônia only [9–10]. In none of our field studies in this region of the Triângulo Mineiro and Alto Paranaiba have we found bats of the genus *Pteronotus* spp. These bats are found in Brazil, in the Amazonian, *cerrado* and Atlantic forest biomes, living mainly near water sources, in caves, and under bridges. *Anoura caudifer*, in which *T. wauwau* was detected, is quite common in Brazil, and can be found in all biomes, particularly in humid forests and in areas with both primary and secondary vegetation.

*T. wauwau* is phylogenetically associated with Australian trypanosomes, possibly constituting one more piece of evidence that *T. cruzi* may have evolved from the recent dispersion of an ancestral bat host across several continents [9]. The finding of bats positive for a disease at aplace where that disease is not endemic, rather is introduced from neighboring areas, demonstrates the considerable mobility of these animals, often involving migrations over long distances, including to urban areas, thereby acting as potential agents of zoonoses.

## References

1. Molyneux DH. Trypanosomes of bats. In: Kreier JP., Baker JR, editors. Parasitic Protozoa. New York: Academic Press; 1991. pp. 95–223.

2. Thomas ME, Rasweiler JJ IV, D’Alessandro A. Experimental transmission of the parasitic flagellates Trypanosoma cruzi and Trypanosoma rangeli between triatomine bugs or mice and captive neotropical bats. Mem Inst Oswaldo Cruz. 2007; 102: 559–565.

3. Gómez-Hernández C, Bento EC, Rezende-Oliveira K, Nascentes GAN, Barbosa CG, Batista LR, et al. Leishmania infection in bats from a non-endemic region of leishmaniasis in Brazil. Parasitol. 2017; 144: 1980–1986.

4. Bento EC, Gómez-Hernández C, Batista LR, Anversa L, Pedrosa AL, Lages-Silva E, et al. Identification of bat trypanosomes from Minas Gerais state, Brazil, based on 18S rDNA and Cathepsin-L-like targets. Parasitol Res. 2018; Forthcoming.

5. Ramirez LE, Machado MI, Maywald PG, Matos A, Chiari E, Silva EL. Primeira evidência de Trypanosoma rangeli no sudeste do Brasil, região endêmica para doença de Chagas. Rev Soc Bras Med Trop. 1998; 31: 99–102.

6. Maia da Silva F, Marcili A, Lima L, Cavazzana M Jr., Ortiz PA, Campaner M, et al. Trypanosoma rangeli isolates of bats from Central Brazil: genotyping and phylogenetic analysis enable description of a new lineage using spliced-leader gene sequences. Acta Trop. 2009; 109: 199–207.

7. Ramírez JD, Tapia-Calle G, Muñoz-Cruz G, Poveda C, Rendón LM, Hincapié E, et al. Trypanosome species in neo-tropical bats: Biological, evolutionary and epidemiological implications. Infect Genet Evol. 2014; 22: 250–256. https://doi.org/10.1016/j.meegid.2013.06.022

8. Pinto CM, Ocaña-Mayorga S, Tapia EE, Lobos SE, Zurita AP, Aguirre-Villacís F, et al. Bats, trypanosomes, and triatomines in Ecuador: New insights into the diversity, transmission, and origins of Trypanosoma cruzi and Chagas disease. PLoS ONE. 2015; 10: e0139999. https://doi.org/10.1371/journal.pone.0139999

9. Lima L, Espinosa-Alvarez O, Pinto CM, Cavazzana M, Pavan AC, Carranza JC, et al. New insights into the evolution of the Trypanosoma cruzi clade provided by a new trypanosome species tightly linked to Neotropical Pteronotus bats and related to an Australian lineage of trypanosomes. Parasit Vectors. 2015; 8: 657.

10. Da Costa AP, Nunes PH, Leite BHS., Ferreira JIGDS, Tonhosolo R, da Rosa AR, et al. Diversity of bats trypanosomes in hydroeletric area of Belo Monte in Brazilian Amazonia. Acta Trop. 2016; 164: 185–193. https://doi.org/10.1016/j.actatropica.2016.08.033

11. De Lima H, Rodrigues N, Barrios MA, Avila A, Canizles I, Gutierrez S. Isolation and molecular identification of Leishmania chagasi from a bat (Carollia perspicillata) in northeastern Venezuela. Mem Inst Oswaldo Cruz. 2008; 103: 412–414.

12. Berzunza-Cruz M, Rodriguez-Moreno A, Gutierrez-Granados G, Gonzalez-Salazar C, Stephens CR, Hidalgo-Mihart M, et al. Leishmania (L.) mexicana infected bats in Mexico: Novel potential reservoirs. PLoS Negl Trop Dis. 2015; 9: e0003690. https://doi.org/10.1371/journal.pntd.0003690

13. Shapiro JT, Lima Junior MSC, Dorval MEC, França AO, Cepa Matos MFC, Bordignona MO. First record of Leishmania braziliensis presence detected in bats, Mato Grosso do Sul, southwest Brazil. Acta Trop. 2013; 128: 171–174.

14. De Oliveira FM, Costa LH, De Barros TL, Ito PK, Colombo FA, De Carvalho C, et al. First detection of Leishmania spp. DNA in Brazilian bats captured strictly in urban areas. Acta Trop. 2015; 150: 176–181.

15. De Castro Ferreira E, Pereira AAS, Silveira M, Margonari C, Marcon GEB, De Oliveira Franca A, et al. Leishmania (V.) braziliensis infecting bats from Pantanal wetland, Brazil: First records for Platyrrhinus lineatus and Artibeus planirostris. Acta Trop. 2017; 172: 217–222.

16. De Rezende MB, Herrera HM, Carvalho CME, Carvalho Anjos EA, Ramos CAN, De Araújo FR, et al. Detection of Leishmania spp. in bats from an area of Brazil endemic for visceral leishmaniasis. Transbound Emerg Dis. 2017; 64: e36–e42. https://doi.org/10.1111/tbed.12597

17. Gregorin R, Taddei VA. Chave artificial para a identificação de molossídeos brasileiros (Mammalia, Chiroptera). J Neotrop Mammal. 2002; 9: 13–32.

18. Vizotto LD, Taddei VA. Chave para a determinação de quirópteros brasileiros., vol 1. Boletim de Ciências, São José de Rio Preto. 1973; pp. 1–72.

19. Reis NR, Shibatta OA, Peracchi AL, Pedro WA, Lima IP. Sobre os morcegos brasileiros. In: Reis NR, Peracchi AL, Pedro WA, Lima IP, editors. Morcegos do Brasil, Londrina; 2007. pp. 17–25.

20. Garcia L, Kindt A, Bermudez H, Llanos-Cuentas A, De Doncker S, Arevalo J, et al. Culture-independent species typing of neotropical Leishmania for clinical validation of a PCR-based assay targeting heat shock protein 70 genes. J Clin Microbiol. 2004; 42: 2294–2297

21. Smith A, Clark P, Averis S, Lymbery AJ, Wayne AF, Morris KD, et al. Trypanosomes in a declining species of threatened Australian marsupial, the brush-tailed bettong Bettongiapenicillata (Marsupialia: Potoroidae). Parasitol. 2008; 135: 1329–1335.

22. Kumar S, Stecher G, Tamura K. MEGA7: Molecular Evolutionary Genetics Analysis version 7.0. Molecular Biology and Evolution (submitted). 2015.

23. TIBCO Software Inc. Statistica (data analysis software system), version 13. 2017. http://statistica.io.

24. Marinkelle CJ, Duarte RCA. Trypanosoma pifanoi n. sp. from Colombian Bats. J Protozool. 1968;15: 621–627.

25. Barnabé C, Brisse S, Tibayrenc M. Phylogenetic diversity of bat trypanosomes of subgenus Schizotrypanum based on multilocus enzyme electrophoresis, random amplified polymorphic DNA, and cytochrome b nucleotide sequence analyses. Inf Gen Evol. 2003; 2: 201–208.

26. Hodo CL, Goodwin CC, Mayes BC, Mariscal JA, Waldrup KA, Hamer SA. Trypanosome species, including Trypanosoma cruzi, in sylvatic and peridomestic bats of Texas, USA. Acta Trop. 2016; 164: 259–266.

27. Szpeiter BB, Ferreira JIGS, Assis FFV, Stelmachtchuk FN, Peixoto Junior KC, Ajzenberg D, et al. Bat trypanosomes from Tapajós-Arapiuns Extractive Reserve in Brazilian Amazon. Rev Bras Parasitol Vet. 2017; 26: 152–158.

28. Cavazzana M Jr., Marcili A, Lima L, Da Silva FM, Junqueira ACV, Veludo HH., et al. Phylogeographical, ecological and biological patterns shown by nuclear (ssrRNA and gGAPDH) and mitochondrial (Cyt b) genes of trypanosomes of the subgenus Schizotrypanum parasitic in Brazilian bats. Int J Parasitol. 2010; 40: 345–355.

29. Ramirez LE, Machado MI, Maywald PG, Matos A; Chiari E; Silva EL. Primeira evidência de *Trypanosoma rangeli* no sudeste do Brasil, região endêmica para doença de Chagas. Revista da Sociedade Brasileira de Medicina Tropical. jan-fev, 1998. 31(1):99–102.

30. Marcili A, Costa AP, Soares HS, Acosta ICL, Lima JTR, Minervino AHH, et al. Isolation and phylogenetic relantionships of bats trypanosomes from different biomes in Mato Grosso, Brazil. J Parasitol. 2013; 99: 1071–1076.

